# Cheese adaptation beyond *Penicillium*: contrasting genomic trajectories in the distantly related *Nectriaceae* molds

**DOI:** 10.64898/2026.10.08.757356

**Authors:** Lily Peck, Jean-Luc Jany, Monika Coton, Océane Savary, Jean-Philippe Vernadet, Alodie Snirc, Tatiana Giraud, Emmanuel Coton, Jeanne Ropars

**Author notes:** Current addresses: (Lily Peck: Department of Ecology and Evolutionary Biology, University of California, Los Angeles, 90095-1438, USA). Equal supervision (Emmanuel Coton, Jeanne Ropars).

## Abstract

Fungal domestication provides a powerful framework for understanding adaptation to human-made environments, yet genomic studies of cheese-associated fungi have largely focused on *Penicillium* molds. Here, we investigated cheese-associated *Nectriaceae*, a phylogenetically distant fungal lineage of ascomycetes, by comparing genomes of nine cheese isolates, belonging to five species, with 22 relatives from other environments. Comparative genomic analyses revealed strikingly contrasting evolutionary trajectories in the two cheese-associated genera. *Longinectria* showed extensive gene family expansions and duplications, particularly affecting nutrient transport, specialised metabolism and fungal recognition, whereas cheese-associated *Bisifusarium* exhibited genome reduction and contraction of many of the same functional systems. In *Bisifusarium domesticum*, these reductions extended to biosynthetic gene clusters, carbohydrate-active enzymes and peptidases, consistent with ecological specialisation. We further identified a 75-kb region with >98% nucleotide identity between two cheese-associated *Bisifusarium* species, strongly supporting recent horizontal transfer. This region contains several genes involved in ion transport and salt tolerance, including functions previously identified in horizontally transferred regions of cheese-associated *Penicillium* populations. Together, our results reveal multiple genomic routes to cheese association and highlight gene-family turnover, adaptive protein evolution and horizontal gene transfer as complementary mechanisms shaping fungal evolution in anthropogenic environments.

## Introduction

Understanding how organisms adapt to novel environments is a central question in evolutionary biology. Domestication is a particularly powerful model for studying adaptation, as it involves recent, strong, and often repeated human-driven selection. Studies ranging from microbes to plants and animals have revealed important genetic and phenotypic changes over relatively short evolutionary timescales (Hufford et al., 2012; Steensels et al., 2019; Warmuth et al., 2012). The genomic processes underlying adaptation have been identified in several domesticated organisms, including horizontal gene transfer, hybridization, gene gain and loss, expansion of gene families, transposable element insertions and regulatory networks modifications (Cao et al., 2026; Domínguez et al., 2020; Schrader and Schmitz, 2019; Steensels et al., 2021).

Moreover, because humans have domesticated phylogenetically distant lineages for similar uses, these domesticated organisms represent ideal models for studying convergent evolution, *i.e.* independent adaptation events of phylogenetically distant lineages to the same ecological niche over short timescales. In fact, phenotypic convergence has been observed in domesticated animals, with either convergent genomic signatures of domestication, involving the same genes, or having occurred through distinct genomic changes (*e.g.* Alberto et al., 2018). Similarly, domesticated crops exhibit multiple convergent phenotypic traits, often involving distinct genomic changes (Cao et al., 2026; Huang et al., 2022), but also sometimes with the same genetic basis, such as for the loss of grain shattering (Jia et al., 2013; Lin et al., 2012).

Domesticated fungi are especially tractable systems for investigating domestication. They have compact genomes, diverse lifestyles and undergo repeated independent transitions into similar human-associated environments (Gladieux et al., 2014; Ropars and Giraud, 2022). Among these, fungi associated with fermented foods, such as cheeses, provide exceptional opportunities to study the genomic mechanisms underlying adaptation to human-made niches that have been produced by humans for thousands of years (Greco et al., 2018; Maixner et al., 2021; McClure et al., 2018; Salque et al., 2013). Because multiple distant fungal lineages have been domesticated for cheese making, they are also excellent models for addressing the question of whether adaptation to human-made environments occurs through the same phenotypes, the same genes and same genomic mechanisms. In *Penicillium* fungi used for cheese making, repeated evolution of traits advantageous for food making has been documented, including faster growth on cheese in *P. camemberti*, *P. biforme* and *P. roqueforti*, and higher tolerance to salt, acidic pH and lactic acid in *P. roqueforti* (Crequer et al., 2023; Ropars et al., 2020). These cheese lineages also displayed strong genetic bottlenecks associated with domestication (Cheeseman et al., 2014; Lo et al., 2023; O’Donnell et al., 2026, 2025; Ropars et al., 2020) and a reduced or complete loss of ability to produce some specialised toxic metabolites, such as andrastin A, mycophenolic acid and PR toxin in *P. roqueforti* (Crequer et al., 2024), and cyclopiazoic acid (CPA) in *P. camemberti* (Ropars et al., 2020). The loss or decrease in toxin production may be beneficial for avoiding food toxicity, or may result from relaxed selection if the fungi are inoculated in cheese at such high quantities that these metabolites are not required against competitors any more. Many of these phenotypic changes have been associated with gains, losses or functional divergence of biosynthetic gene clusters (Crequer et al., 2024; Ye et al., 2024). These studies have thus demonstrated that adaptation to food environments has occurred repeatedly, and often convergently, across distantly related fungal lineages.

Studies on *Penicillium* fungi used for food fermentation, including the cheese fungi *P. roqueforti, P. camemberti* and *P. biforme*, as well as the dry-sausage fungi *P. nalgiovense* and *P. salamii*, have revealed horizontal transfers of hundreds of genes in these domesticated *Penicillium* fungi, that were subsequently recognised as cargo genes carried by giant mobile elements, called *Starships*. These mobile elements, capable of transferring genes between strains and species, are increasingly recognized as important drivers of fungal niche adaptation and domestication (Urquhart et al., 2024). Remarkably, the same *Starships* were identified in independently domesticated and phylogenetically distant lineages, for example between the three cheese molds *P. camemberti*, *P. biforme* and *P. roqueforti* (Cheeseman et al., 2014; O’Donnell et al., 2026; Ropars et al., 2020), as well as between the two dried sausage molds *P. nalgiovense* and *P. salamii* (Lo et al., 2023; O’Donnell et al., 2025). This suggests the repeated transfers of the same genes involved in adaptation.

Beyond horizontal gene transfer, adaptation can also proceed through gene family expansion and contraction, gene gain and loss, and changes in protein-coding sequence driven by natural selection (Borneman et al., 2016; Douglas and Langille, 2019; Gladieux et al., 2014; Ropars et al., 2015). Gene family expansions and duplications increase gene copy numbers, and some copies can subsequently evolve new or specialised functions. In contrast, gene family contraction can reflect the loss of functions that are no longer useful in a novel environment (Kondrashov, 2012). Positive selection can further refine protein function by favouring amino-acid substitutions that improve performance under new ecological conditions (Gladieux et al., 2014). These complementary processes can facilitate adaptation to new ecological niches; yet, compared with horizontal transfers, much less is known about their contribution to adaptation in domesticated cheese fungi.

Other fungal lineages are used for cheese making but have been much less studied than *Penicillium* molds. In particular, the *Bisifusarium domesticum* mold, commercially used for cheese maturation, has been studied from a physiological and metabolic perspective (Savary et al., 2023, 2022), but has not yet been investigated at the genomic level. This species has solely been found in cheese to date, and is valued for its *anticollanti* (anti-stickiness) properties, helping to prevent cheese surface stickiness (e.g. on Raclette, Reblochon) (Bachmann et al., 2005). It belongs to the *Nectriaceae*, a particularly interesting fungal family for studying domestication in cheese, because it represents an evolutionary lineage distant from the well-studied *Penicillium* cheese molds. *Bisifusarium* belong to the Sordariomycetes, whereas *Penicillium* belongs to the Eurotiomycetes, meaning that their adaptation to the same human-made cheese environment provides an excellent system for investigating convergent adaptation and independent domestication across distantly related fungi. The taxonomy of *B. domesticum* has changed several times, from *Trichothecium domesticum* to *Fusarium domesticum* (Bachmann et al., 2005), and more recently to a new genus named *Bisifusarium* (Lombard et al., 2015). The *Nectriaceae* includes around 55 genera, including the well-known *Fusarium genus,* containing important plant pathogens and species capable of producing mycotoxins, making it especially interesting to investigate how adaptation to cheese may have altered traits related to metabolism, competition, and toxin production. Recently, four new Nectriaceae cheese-associated species were discovered through microbiota isolation from 56 cheese rinds (Savary, 2021) and phylogenetic analyses based on 10 genes. These four novel species included two assigned to a novel genus, *Longinectria,* which is not a sister group to *Bisifusarium.* These two species were named *Longinectria lagenoides* and *Longinectria verticilliformis*. Two new sister species belonging to the genus *Bisifusarium* were also described: *Bisifusarium allantoides* and *Bisifusarium penicilloides* (Savary, 2021). These fungi are well suited to the cheese-ripening environments as they grow at low temperatures, tolerate cheese-associated abiotic conditions, and produce volatile compounds that contribute to cheese aroma (Savary et al., 2023, 2022). However, whether they have actually adapted to cheese through genetic changes compared to their wild counterparts has not yet been explored. Here, we therefore addressed the following questions: Have these cheese lineages acquired genes potentially beneficial for cheese making? Do these lineages show evidence of recent horizontal gene transfers, gene family expansion or contraction? Have the biosynthetic gene clusters of secondary metabolites, of CAZymes (carbohydrate-active enzymes) and peptidase (protein-degrading enzymes) been lost or in contrast expanded? Does *B. domesticum*, inoculated for cheese-making, show footprints of bottlenecks, i.e. very low genetic diversity?

To address these questions here, we acquired the genomes of nine cheese-associated *Nectriaceae* species and compared them with 22 available genomes of other *Nectriaceae* species, thriving in various non-cheese habitats. The cheese-associated *Nectriaceae* fungi comprise three groups: the two *Longinectria* species which were recently identified and only found in cheese to date, *B. domesticum*, with which cheese is inoculated, and the two newly identified *Bisifusarium* species, *B. allantoides* and *B. penicillioides*, this later species having been isolated from cheese and other environments. With these genomes, we 1) reconstructed the evolutionary relationships between cheese-associated lineages, and with other *Nectriaceae* species that thrive in non-cheese environments, 2) identified genomic footprints of domestication, *i.e.* gene gains and losses, expansion and reduction of some key gene families, 3) looked for footprints of horizontal gene transfers between cheese strains, 4) investigated the genomic evolution of biosynthetic gene clusters of secondary metabolites, CAZymes and peptidases, and 5) assessed whether species having independently adapted to cheese show convergent genomic changes.

## Material and methods

### Fungal strains

All selected cheese-associated Nectriaceae strains (i.e. belonging to the *Bisifusarium allantoides*, *Bisifusarium domesticum, Bisifusarium penicilloides, Longinectria lagenoides* or *Longinectria verticilliformis* species) (Table 1) were obtained from previous studies (Savary, 2021) and preserved in either the ESE or UBO Culture collection (https://www.univ-brest.fr/ubocc/fr). All specimens studied originated from monosporal cultures.

**Table 1.** Origin and genome quality statistics for *Nectriaceae* strains isolated from cheese and sequenced in this study. Although ESE00026 (*Fusarium solani)* was isolated from a cheese, this species is not considered as cheese-associated but rather a contaminant.

| Strain name | Species (Lombard) | Strain origin | BUSCO score | Genome size (Mb) | N50 (kb) | # gene | GC content (%) |
| --- | --- | --- | --- | --- | --- | --- | --- |
| UBOCC-A-120036 | *Bisifusarium allantoides* | Soft cheese type | 99.2 | 31.1 | 525.7 | 10027 | 49.57 |
| UBOCC-A-120021 | *Bisifusarium penicilloides* | Cheese (Mont d'Or) | 99.1 | 31.6 | 509.1 | 9647 | 47.4 |
| ESE00042 | *Bisifusarium domesticum* | Cheese (Eguia) | 99.1 | 34.2 | 140.8 | 9459 | 48.21 |
| ESE00064 | *Bisifusarium domesticum* | Cheese (Bethmale, tomme pyrénées) | 99.1 | 34.1 | 120.4 | 9415 | 48.24 |
| LMA587 | *Bisifusarium domesticum* | Artisanal cheese with wash crust, Quebec Canada | 99.1 | 33.5 | 125.0 | 9354 | 48.36 |
| UBOCC-A-120004 | *Bisifusarium domesticum* | Commercial adjunct culture | 99.2 | 33.4 | 53.0 | 9289 | 48.43 |
| ESE00026 | *Fusarium solani* | Contaminated cheese (La Toudeille) | 99.7 | 49.3 | 432.5 | 15967 | 50.42 |
| ESE00140 | *Longinectria lagenoides* | Cheese (Tilsit) | 99.5 | 59.6 | 99.2 | 15955 | 50.09 |
| mUBOCC-A-120039 | *Longinectria lagenoides* | Cheese (Vacherin Fribourgeois) | 99.3 | 59.7 | 100.4 | 16081 | 49.96 |
| UBOCC-A-120043 | *Longinectria verticilliforme* | Cheese (Alpeggio) | 99.8 | 48.1 | 157.3 | 13742 | 50.42 |

### Culture and DNA extraction

For genome sequencing, ten strains (Table 1) were grown for seven to ten days on malt extract agar at 25°C and genomic DNA was extracted using the Nucleospin Soil Kit (Macherey-Nagel, Duren, Germany). For all DNA extractions, quantity and quality were determined by Nanodrop 2000 (Thermofisher Scientific, Waltham, US) and by the Qubit 4 fluorometer (Thermofisher Scientific, Waltham, US) combined with the Qubit dsDNA BR Assay Kit to estimate if the requirements were reached for whole genome sequencing.

### Genome sequencing, short-read assemblies and annotation

Short-read sequencing was performed with Illumina HiSeq 2500 paired-end technology (Illumina Inc.), with an average insert size of 400 bp at the GenoToul GeT INRAe platform (Genopole de Toulouse, France). Illumina reads were trimmed and adapters cleaned with Trimmomatic v0.39 (Bolger et al., 2014). Leading or trailing low-quality or base pairs below a quality score of three were removed. For each read, only parts that had an average quality score higher than 20 on a four base pair (bp) window are kept. After these steps, only reads with a length of at least 36 bp were kept. Cleaned Illumina reads were assembled with SPAdes v3.15.5 (Prjibelski et al., 2020) not using unpaired reads with “--careful” parameter.

All genomes, including those already published, were automatically annotated to have a homogenised dataset. We used Braker (v3.0.8) (--fungus --gff3) (Gabriel et al., 2024) for genome structure annotation using the fungi_odb10 BUSCO dataset (Manni et al., 2021). Four more rounds were then rerun with Braker using the previous runs of augustus protein hints. To assess the quality of the structural annotation, the resulting protein dataset was then analyzed using the fungi_odb10 BUSCO dataset and functionally annotated using Funannotate (v1.8.15) (Palmer & Stajich, 2020). We generated a funannotate database with the “setup” command (-b fungi). Next, the funannotate iprscan wrapper (interproscan-5.56-89.0) (Jones et al., 2014) was used to annotate protein domains and eggnog-mapper (v2.1.12) (Cantalapiedra et al., 2021) for Eggnog functions. Functional and structural annotations were finally aggregated with the “annotate” function.

### Orthology inference and species tree

We used protein coding sequences from the 32 published genomes belonging to the *Nectriaceae* included in this study (Table S1). First, we cleaned the input protein sequences to remove all transcripts except the longest isoform using ‘primary_transcript.py’ (https://github.com/ davidemms/OrthoFinder). We then used OrthoFinder v3.0.1b1 (Emms et al., 2025; Emms and Kelly, 2019, 2015) to assemble orthogroups of protein coding genes. OrthoFinder inferred orthogroups and their corresponding gene trees by clustering together protein sequences based on similarity. These gene trees were summarised to build a rooted species tree which was used to characterise gene duplication events using gene trees together with the species tree. OrthoFinder provides a list of single-copy orthologues in all species considered: we used these genes with a one-to-one relationship to build a “supermatrix” species tree and to test for selection (described below).

The species tree was inferred from single-copy orthologues identified by OrthoFinder. Of the 3,656 single-copy orthogroups, we retained the 3,258 whose coding sequences matched their proteins in all genomes. Proteins were aligned with MAFFT v7.526 L-INS-i ‘--localpair --maxiterate 1000’ (Katoh and Standley, 2013) and trimmed with trimAl v1.5.1 ‘-automated1’ (Capella-Gutiérrez et al., 2009). We mapped each coding sequence onto its protein alignment, and discarded orthogroups with trimmed alignments shorter than 100 amino acids, leaving 3,201 genes. The best-fitting substitution model for each protein alignment was selected with ModelFinder under BIC ‘-m MF’ (Kalyaanamoorthy et al., 2017), before the alignments were concatenated into a supermatrix of amino acid sites, partitioned by gene, with each partition assigned its ModelFinder model. A maximum-likelihood tree was inferred using IQ-TREE v3.1.3 (Wong et al., 2026) from the protein supermatrix, with per-partition substitution models as selected by ModelFinder (-m MF) and 1,000 ultrafast bootstrap replicates (-B 1000) with the ‘--bnni’ correction (UFBoot2, (Hoang et al., 2018). The final tree was rooted with *Xenoacrimonium recifei* based on predetermined phylogenetic relationships..

Following selection of the final rooted species tree, OrthoFinder was re-run from the existing gene trees using the rooted coding sequence species tree, ensuring that orthology relationships were inferred consistently with the *X. recifei* root. Gene-family analyses were based on the root level hierarchical orthogroups from this OrthoFinder re-run.

### Gene family expansions and contractions

First, we transformed the rooted species tree into an ultrametric tree with penalised-likelihood rate smoothing in ‘chronos’ (ape v5.8, Paradis and Schliep, 2019), with a correlated-rates model and the default smoothing parameter (λ = 1). In the absence of fossil or absolute-time calibrations, the root age was fixed to 1. All branch lengths were subsequently multiplied by 100 to obtain a tree depth of 100 relative time units, preserving the proportional spacing of divergence events and resolving non-finite starting likelihoods encountered in CAFE5 at a tree depth of 1.

Gene family expansions and contractions were analysed using gene counts obtained from root-level hierarchical orthogroups in OrthoFinder (HOGs, ‘N0.tsv’). CAFE v5.1.0 models changes in gene family size across a phylogeny as a stochastic birth-death process (Mendes et al., 2020). It estimates λ, the per-gene rate of gene birth and death per unit time (De Bie et al., 2006; Hahn et al., 2005), and uses it to compute the likelihood of the observed family sizes and to infer ancestral family sizes. We used the same ultrametric tree in CAFE as we did to make the species tree (described above). We retained gene families that were present on both sides of the root, that is, in *X. recifei* and in at least one other genome, to meet the model assumption that analysed families were present in the common ancestor of all sampled taxa, leaving 11,878 gene families (7,141 excluded). No retained family contained >100 copies in any genome, and all retained families were analysed together. A single gene birth-death rate (λ), shared across branches and families, was estimated jointly with a global gene-count error model to account for potential assembly errors. Three independent runs yielded essentially identical estimates (-lnL = 276,541; λ = 0.00317 per relative time unit; error rate = 0.028). A family was considered to have changed significantly if its size varied more than expected under the fitted model across the tree using a family-wide *P* < 0.05. For each node, we counted these families as expanded or contracted when their inferred ancestral size increased or decreased on the branch leading to that node.

### Functional enrichment

Functional enrichment was assessed separately for gene-family sets identified by CAFE5 and genes showing evidence of positive selection, using orthogroups as the unit of analysis. CAFE5 sets comprised significantly expanded families on the ancestral *Longinectria* branch and all families reconstructed as contracting on the *B. domesticum* branch. Annotations included Pfam domains, InterPro annotations, eggNOG assignments, COG functional categories, gene ontology terms and predicted product descriptions. For families absent from the focal lineage, annotations were obtained from orthologues retained in related taxa. Each focal set was compared with its corresponding analysis background using two-sided Fisher’s exact tests, performed separately for each annotation type. Only terms represented in at least 30 orthogroups within the relevant background were tested. *P*-values were adjusted using the Benjamini-Hochberg procedure within each focal set and annotation type; terms with adjusted *P* < 0.05 and higher frequencies in the focal set were considered enriched.

### Starship identification

The tool *Starfish* (Gluck-Thaler, 2023) was used to identify *Starships* and their captains, that encode DUF3435-domain proteins providing the putative mobility function. The *Gene finder* and *Element finder* modules were launched. Despite some captains having been identified in the genomes (Table S1), no mobile elements were found with the Element finder module. This was however expected as our genome assemblies exhibited rather low quality. We thus use BLAST (blastn) to identify regions >5kb and with a nucleotide identity > 98% between cheese species to detect horizontal transferred regions.

*Biosynthetic gene clusters, carbohydrate-active enzymes and peptidases* We only included cheese species from *Bisifusarium* and *Longinectria* genera and the wild relative of *Bisifusarium* cheese species *B. dimerum* for this analysis because an assembly was publicly available. Raw sequencing reads or genome assemblies were processed using the FunFlux v1.0.4 pipeline, a Snakemake-based workflow designed for the assembly and annotation of fungal genomes from Illumina short-read data (Antonielli, 2026). Biosynthetic gene clusters were predicted using antiSMASH v8.0 (Blin et al., 2025) integrated within the FunFlux pipeline. Predicted biosynthetic gene clusters were subsequently grouped into gene cluster families using BiG-SCAPE (Navarro-Muñoz et al., 2020) with a similarity cut-off of 0.4. Presence/absence matrices of gene cluster families across genomes were generated in R (v4.6.0) and visualized as heatmaps using the *pheatmap* package. Comparative analyses of carbohydrate-active enzyme (CAZyme) and peptidase repertoires were performed in R (v4.6.0) using the annotation files generated by the FunFlux pipeline. Within FunFlux, CAZymes were annotated using dbCAN (Zheng et al., 2023) based on the CAZy database, whereas proteases and protease inhibitors were identified using the microbial eukaryote reference oriented peptidase (MEROPS) database (Rawlings et al., 2014). Abundance matrices were generated at both the family and functional class levels for CAZymes (GH, glycoside hydrolases; GT, glycosyltransferases; CE, carbohydrate esterases; AA, auxiliary activities; CBM, carbohydrate binding molecules; and PL, pectate lyases) and peptidases (Aspartic, Cysteine, Metallo, Serine, Threonine, Mixed, and Unknown). Heatmaps of detailed family repertoires were generated from protein counts standardized independently for each family across genomes (Z-scores), allowing the visualization of relative enrichments and depletions irrespective of absolute family size. Principal component analyses were performed on centered and scaled abundance matrices to compare the functional profiles of the different genomes.

## Results

### Comparative genomics resolves *Bisifusarium* and *Longinectria* lineages and identifies two *B. domesticum* commercial cultures

Genome-wide analyses of cheese-associated *Bisifusarium* and *Longinectria* resolved their phylogenetic relationships within the *Nectriaceae* and revealed remarkably low genetic diversity among *B. domesticum* isolates, consistent with the presence of distinct clonal cultures and strong bottlenecks. We generated short-read genome assemblies for ten strains isolated from different cheese rinds, encompassing two *L. lagenoides* strains, *L. verticilliforme, B. allantoides, B. penicillioides,* and four *B. domesticum* strains (Table 1). We also generated a new short-read genome assembly of a *Fusarium solani* strain isolated from a contaminated cheese. Among the species sampled, *B. domesticum* was the only known species exclusively isolated from cheese to date and is intentionally added during cheese production. Other members of *Bisifusarium* occur in a range of environments beyond cheese. To our knowledge, the two cheese-associated species *L. lagenoides* and *L. verticilliforme* are the only known members of this genus. We compared these nine newly sequenced genomes with published genomes from one additional *B. domesticum* strain and 22 non-cheese species within the same *Nectriaceae* family (Table S1). Our dataset thus included 32 haploid genomes.

We identified 28,105 orthologous groups of protein-coding genes, of which 4,752 orthogroups contained at least one gene copy from all 32 genomes, while there were 3,656 single-copy gene orthogroups. A species tree reconstructed from single-copy gene trees supported the phylogenetic relationships established in earlier studies with fewer genetic loci (Savary, 2021). Each of *Bisifusarium* and *Longinectria* forms a monophyletic clade within the Nectriaceae family, both with 100% bootstrap branch support in the species tree (Figure 1A, 1000 replicates). Average nucleotide identity dropped below 75% between the *Longinectria* and *Bisifusarium* genera (Figure 1B, Table S2). Within these two genera, average nucleotide identity between the different species was >85% (Table S2).

**Figure 1.**
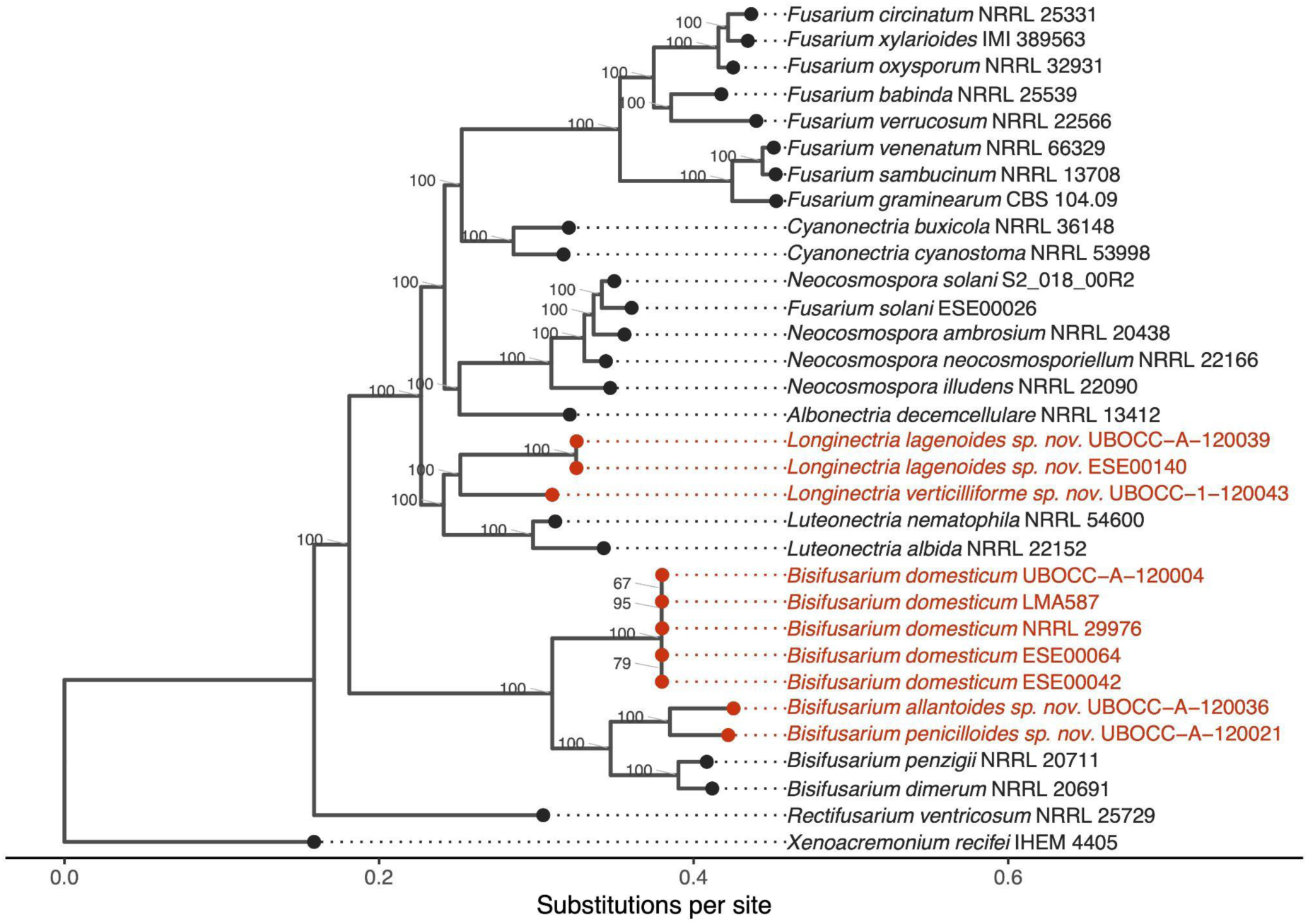
Phylogeny across the *Nectriaceae* family. **A** Species tree based on 3,201 single-copy shared orthogroups. Annotated branches indicate bootstrap branch support values (1000 replicates). Red tip labels indicate the cheese-associated *Longinectria* and *Bisifusarium* species, which were all sequenced in this study except *B. domesticum* NRRL 29976. Full strain details on the studied strains are given in Tables 1 and S1.

Comparison among the five *B. domesticum* genomes indicated the existence of two clonal commercial cultures. We used the four genomes acquired here together with the publicly available NRRL 29976 assembly to assess the genetic diversity within the only species recovered in this study with a known role in cheese-making, i.e, *B. domesticum*. As no high-quality genome assembly of *B. domesticum* was available, we mapped short raw reads of these five *B. domesticum* strains against *B. domesticum* ESE00042, as it was the one combining the best BUSCO score (99.1), the best N50 and the least fragmented assembly. Fewer than 400 SNPs were identified across *B. domesticum* strains, except for UBOCC-A-120004, a known commercial adjunct culture, which displayed 7,150 SNPs (Table S3). The four other isolates, with few genetic differences between each other, may thus belong to a clonal lineage, likely used to inoculate cheese, as another commercial culture, closely related but distinct from UBOCC-A-120004. Mapping all raw reads against UBOCC-A-12004 confirmed that this strain was different from all others, with about 8,500 SNPs identified for each mapped genome (Table S3). Given the genome size of *B. domesticum* (33.5 Mb), this represents a 0.026% difference between the known commercial strain and the other four isolates.

### Shared and lineage-specific signatures of adaptation to cheese

Comparative genomic analyses revealed contrasting patterns of gene family evolution in the two independently cheese-associated lineages, *Bisifusarium* and *Longinectria.* Gene family turnover was extensive throughout the sampled *Nectriaceae,* with both expansions and contractions occurring repeatedly across the phylogeny. Within *Longinectria*, gene family expansions and duplications predominated. In contrast, contractions predominated in *Bisifusarium,* and cheese-associated species had particularly compact genomes (median 33.4 Mb) and low predicted gene numbers (median 9,415). Functional analyses identified changes in nutrient transport and metabolism, secondary metabolism, and carbohydrate-active enzyme and peptidase repertoires.

### Gene family expansion and duplication in the Longinectria genus

Both cheese-associated strains in the *L. lagenoides* lineage displayed gene family expansions and gene duplications, affecting metabolic, transport and extracellular enzyme repertoires. Indeed, we detected substantial heterogeneity in rates of gene family expansion and contraction across the phylogeny (Figure 2). Gene family changes in *Longinectria* shifted from predominantly contractions at the genus branch to predominantly expansions on the *L. lagenoides* branch. Along the *L. lagenoides* branch, 538 gene families were inferred to have expanded and 377 contracted, representing the largest expansion in the phylogeny. This expansion signal was supported by an independent analysis of gene duplications, whereby the *L. lagenoides* branch contained 554 well-supported duplication events, substantially more than elsewhere in the phylogeny (Figure S1). Although several other lineages, including the plant pathogenic *Fusarium* species, *F. oxysporum* and *F. xylarioides* had 486 expanded gene families, none showed the same combination of extensive expansion and gene duplication. In *L. lagenoides*, gene family expansions were enriched for carbohydrate transport and metabolism, and major facilitator transporters include amino acid permeases (Tables S4 and S5). Other enriched functions included heterokaryon incompatibility, which was also strongly enriched in families containing duplications in *L. lagenoides* (Table S9).

**Figure 2.**
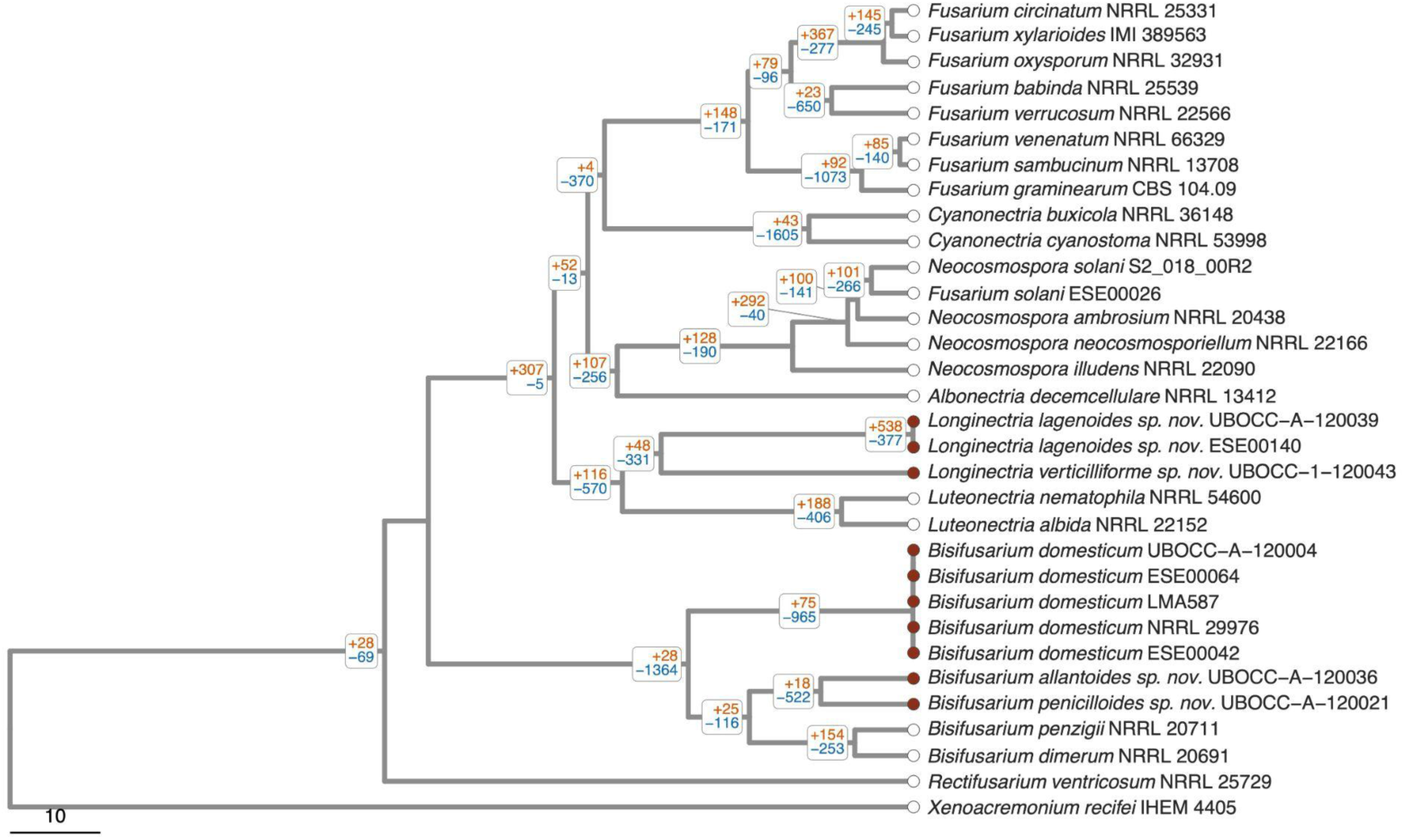
Gene-family expansions and contractions across the *Nectriaceae* family. The phylogeny comprises 32 genomes and is based on the species tree (Figure 1A). Branch lengths were scaled to relative time using chronos, with the root age set to 100; the scale bar represents 10 relative time units. Branch labels indicate the numbers of gene families inferred by CAFE to have expanded (orange, +) and contracted (blue, -) along branches leading to internal nodes. Counts only include families with significant size variation across the (family-wide *P* < 0.05). Filled circles indicate strains isolated from cheese.

These functional expansions also affected extracellular enzyme repertoires. *Longinectria lagenoides* in particular possessed consistently larger carbohydrate active enzyme (CAZyme) repertoires across all major enzyme classes (GH, glycoside hydrolases; GT, glycosyltransferases; CE, carbohydrate esterases; AA, auxiliary activities; CBM, carbohydrate binding molecules; and PL, pectate lyases; Figures S4 and S5) together with substantially larger complements of serine, metallo- and cysteine-peptidases than the cheese-associated *Bisifusarium* species.

### Gene family and functional repertoire contraction in the Bisifusarium genus

The whole *Bisifusarium* lineage, including both cheese- and non-cheese strains, showed extensive gene family contraction and genome reduction, with reduced repertoires related to nutrient transport, specialised metabolism and extracellular degradation. Gene family contractions dominated on both the *Bisifusarium* stem, ancestral to cheese-associated and environmental species, as well as the branch leading to *B. domesticum.* These branches contained 1,364 and 965 contracted families, respectively, compared with 28 and 75 expanded families (Figure 2). While other parts of the phylogeny also showed inferred contractions (including *Cyanonectria* with 1,605 and the clade containing *F. graminearum, F. sambucinum,* and *F. venenatum* with 1,073 contractions), only *Bisifusarium* shows this contraction biased trajectory with few expansions across multiple successive branches.

Families contracted in *B. domesticum* were enriched for carbohydrate transport and metabolism, and amino acid transport and metabolism. Enriched transporter domains included major facilitator transporters and amino acid permeases (Tables S7 and S8). Secondary-metabolite biosynthesis, transport and catabolism was also enriched, encompassing 95 contracted families. Of these, 92 had reconstructed copy numbers of zero at the *B. domesticum* ancestral node and were absent from all five *B. domesticum* annotated gene sets. All 95 families were retained in at least one other sampled *Bisifusarium* species. Other enriched functions among contracted *B. domesticum* families included fungal transcription-factor and protein kinase domains (Tables S7 and S8). Genome reduction in *B. domesticum* thus disproportionately affected families associated with nutrient transport and metabolism, including specialised metabolic functions.

In the genomes of the cheese-associated species, we also found substantial reductions in biosynthetic gene clusters, CAZymes (carbohydrate active enzymes), and peptidases, indicating a broad contraction of the functional repertoire used to acquire nutrients and mediate ecological interactions. The three cheese-associated *Bisifusarium* species (*B. allantoides, B. penicillioides, B. domesticum*) carried fewer biosynthetic gene clusters than the closely related non-cheese *Bisifusarium* and other non-cheese species examined. *Bisifusarium domesticum* had the smallest repertoire, with 18 predicted clusters, compared with up to 37 in its close wild relative *B. dimerum* (Figure S2).

A similar pattern of genome reduction was observed in extracellular enzyme repertoires. Principal component analysis of the combined CAZyme and peptidase repertoires separated the three cheese-associated *Bisifusarium* species from *B. dimerum* and the cheese-associated *Longinectria* (Figure 3). Cheese-associated *Bisifusarium*, particularly *B. domesticum,* had smaller CAZyme (carbohydrate-active enzyme) repertoires across nearly every enzyme class (Figure S4 and S5). In contrast, the wild species *B. dimerum* retained larger repertoires of families associated with plant cell-wall degradation, including cellulose oxidation (AA9), hemicellulose degradation (GH54, GH67 and CE1) and pectin degradation (PL1) degradation. These differences distinguished the cheese-associated species from their environmental relative in functions associated with the utilisation of plant substrates. Peptidase repertoires also differed between the cheese-associated lineages. Cheese-associated *Bisifusarium* encoded fewer serine, metallo-and cysteine-peptidases than the two cheese-associated *Longinectria* species, with *B. domesticum* having the smallest repertoires (Figures S6 and S7). Differences occurred both in total peptidase numbers and in the representation of individual microbial eukaryote reference oriented peptidase (MEROPs) families.

**Figure 3.**
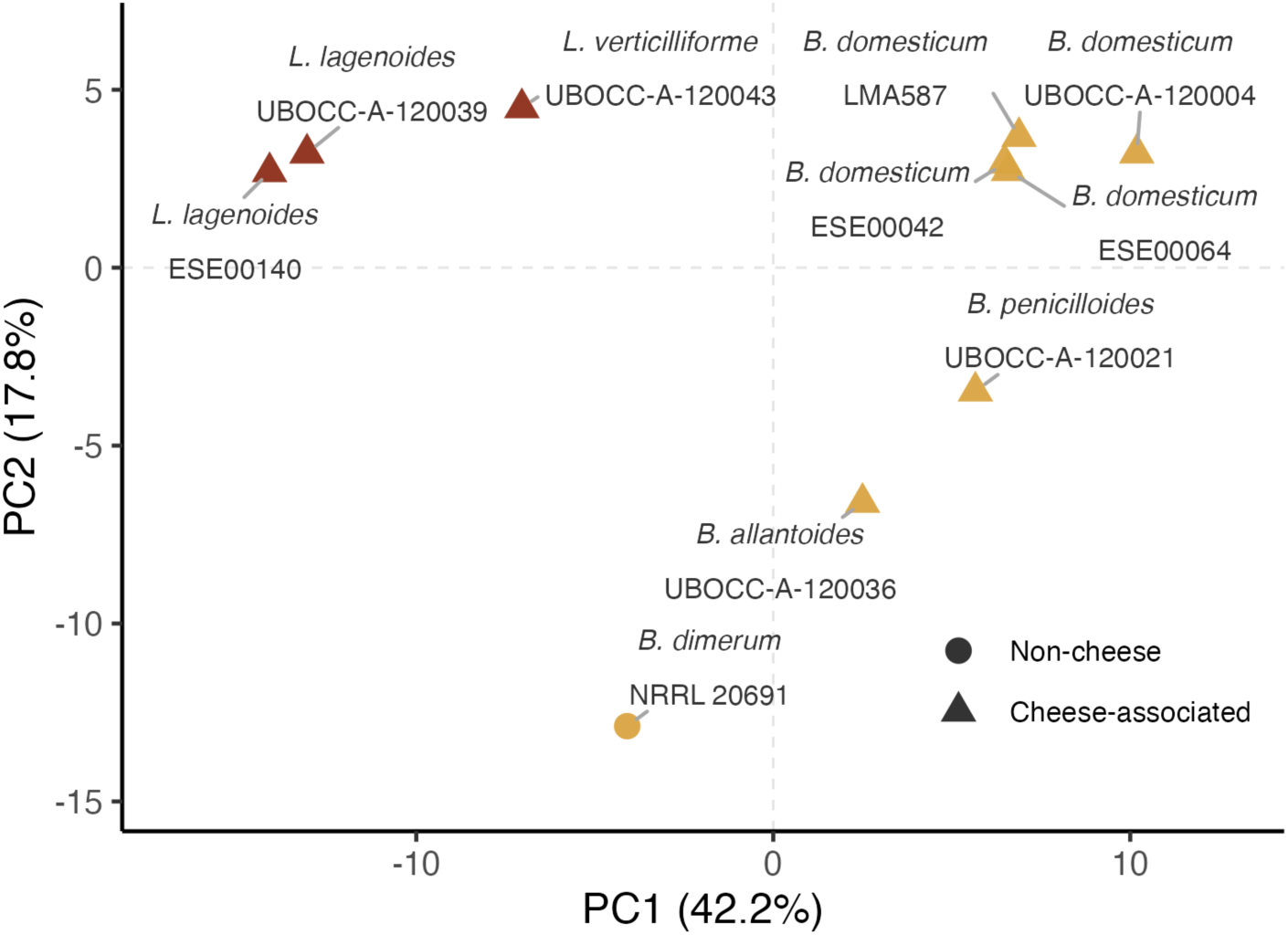
Principal component analysis of carbohydrate active enzymes (CAZymes) and microbial eukaryote reference oriented peptidase (MEROPs) functional profiles across genomes. PCA was performed on centered and scaled counts of CAZymes and MEROPS families. Points represent individual genomes.

Consistent with this long-term reduction in gene content, the cheese-associated *Bisifusarium* species (*B. domesticum, B. allantoides, B. penicillioides)* possessed among the smallest genomes and lowest predicted gene numbers within the dataset (Figure S2; Table S6). The cheese-associated *Longinectria* genomes contained substantially more genes than *Bisifusarium*, although they still encoded fewer genes than would be predicted from their comparatively large genome sizes across the phylogeny. The two non-cheese *Bisifusarium* species (*B. dimerum* and *B. penzigii*) displayed intermediate genome sizes, although these did not differ significantly from those of the cheese-associated species (Wilcoxon test, W = 14, adjusted *P* = 0.13).

### Horizontal transfer of a salt tolerance gene cluster between *Bisifusarium* cheese fungi and potentially numerous starships

We detected numerous *Starship* captains in the genomes studied, encoding DUF3435-domain proteins that are thought to enable the mobility of these giant mobile elements (Table S1). Of the cheese-associated fungi, the two *Longinectria* species had the highest number of captains, with n = 18 in *L. verticilliformis* and n = 11–12 in *L. lagenoides*. In contrast, cheese-associated *Bisifusarium* species had fewer than 10 captains, with only 2 and 3 captains identified in *B. allantoides* and *B. penicillioides*, respectively. We observed a weakly positive correlation between genome size and captain number (Spearman’s rho = 0.45, *P* = 0.0092, Figure S9), suggesting that captain abundance scales broadly with genome size across the Nectriaceae. Interestingly, the reduction in genome size observed in cheese-associated *Bisifusarium* was not accompanied by a disproportionate loss of captains. The genome assemblies were too fragmented (Table 1) to identify complete *Starships*-like regions using *Starfish* (Gluck-Thaler, 2023).

We identified a 75 kb region between *B. penicillioides* and *B. allantoides* with >98% sequence identity (Figure 4), while the average nucleotide identity (ANI) was <84% between the two species (Table S2). Such near 100% sequence identity across the entire 75 kb region, including non-coding sequences, as well as the complete absence of synteny of flanking regions between the two genomes, strongly suggests horizontal transfer. Furthermore, several genes within this putative horizontally transferred region encoded functions previously identified in the salt tolerance cluster of a *Starship-*like region in *Penicillium* cheese fungi (O’Donnell et al., 2025), including ENA2, a central component involved in Na+ efflux to allow salt tolerance, TRK involved in potassium transport, two chromate transporters, a phosphate transporter and a sodium symporter. In addition, at one edge of this region we identified a SANT/Myb gene, previously found in *Starship*-like regions in cheese-associated *Penicillium* fungi (O’Donnell et al., 2025). We could not identify any captain in this region, but both genome assemblies were highly fragmented. We did not identify any other regions with >98% nucleotide identity between the two cheese-associated genera, between the two *Longinectria* species, or between *B. domesticum* and the other two cheese-associated *Bisifusarium* species.

**Figure 4.**
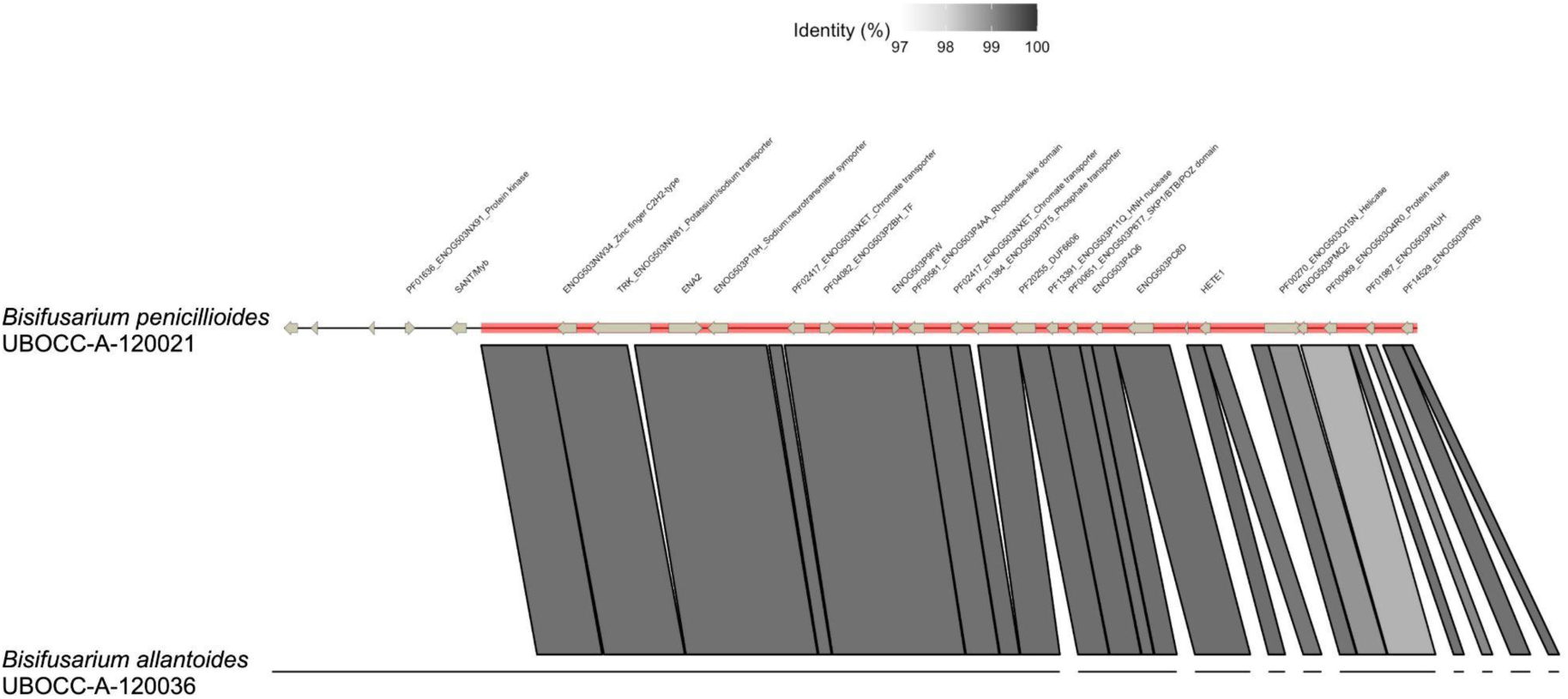
Schematic representation of the putative horizontal transfer between *Bisifusarium allantoides* UBOCC-A-120036 and *B. penicillioides* UBOCC-A-120021. The region sharing a nucleotide identity >98% is represented in red on UBOCC-A-120021, location of the genes are indicated by grey arrows, with their putative functions above. The absence of blocks between the two genomic sequences on either side of the putative transfer indicate the absence of synteny between the two genomes.

## Discussion

Cheese represents a highly specialised, human-created environment characterised by high salt concentrations, abundant protein and lipid substrates, and reduced environmental variability compared to natural habitats. The molds used for cheese making studied so far exhibit footprints of adaptation to cheese, in terms of genetic and phenotypic changes compared to their wild counterparts. For example, phylogenetically distant *Penicillium* cheese fungi have acquired the same salt tolerance gene cluster via *Starships* (O’Donnell et al., 2025). Several other *Starships* carrying genes beneficial for cheesemaking have been identified in distantly related *Penicillium* cheese fungi, including some involved in antagonistic interactions or lactose metabolism (Cheeseman et al., 2014; O’Donnell et al., 2026, 2025; Ropars et al., 2020, 2015). In the yeast *Geotrichum candidum,* naturally present in raw milk and intentionally used in some cheese productions, cheese-associated populations lack two beta-lactamase-like genes involved in xenobiotic clearance, likely due to relaxed selection (Bennetot et al., 2023). Similarly, in *Saccharomyces cerevisiae*, domesticated populations show extensive gene loss and pseudogenisation, with cheese-associated strains among those exhibiting the greatest loss of genomic regions (Legras et al., 2018).

The present study reveals that two phylogenetically distant fungal lineages within the Nectriaceae family, *Bisifusarium* and *Longinectria,* followed strikingly different evolutionary trajectories in adapting to cheese. Indeed, *Longinectria* species had larger genome sizes and had primarily expanded their gene repertoire through the massive expansion of some gene families potentially beneficial for cheese production. In contrast, *Bisifusarium* species had smaller genome sizes and underwent a drastic loss and contraction of gene families no longer necessary in this ecological niche.

In *L. lagenoides,* to date only isolated from cheese, expanded families were enriched for nutrient transport, transcriptional regulation, carbohydrate and secondary metabolism. Several studies have linked nutrient transport and carbohydrate metabolism with adaptation to dairy environments, including in *Penicillium* (Ropars et al., 2015) and *Kluyveromyces* fungi (Varela et al., 2019). Changes in secondary metabolism have also accompanied cheese domestication in *Penicillium,* whereby experimental evolution reduced secondary metabolite production and downregulation of associated pathways (Bodinaku et al., 2019). Expansion and duplication analyses both identified strong enrichment of HET-domain families associated with fungal recognition and incompatibility. In contrast, cheese-associated *Bisifusarium* species exhibited widespread contraction of many of these functional systems.

Contrasting patterns between cheese-associated *Longinectria* and *Bisifusarium* species were also observed in terms of biosynthetic gene clusters, carbohydrate-active enzymes (CAZymes) and peptidases. Indeed, the three cheese-associated *Bisifusarium* species, and particularly *B. domesticum*, inoculated in cheese by producers, exhibited substantial reductions in secondary metabolite biosynthesis, carbohydrate degradation and extracellular proteolysis. In contrast, *Longinectria* retained comparatively large and diverse repertoires across these functional categories. Comparative genomic studies have linked variation in these functional repertoires to fungal nutritional strategies and lifestyles (Dort et al., 2023; Zhao et al., 2013). The contrasting CAZyme repertoires involved particular functional groups rather than a uniform difference across all carbohydrate-active enzymes. *Longinectria* species, and particularly *L. lagenoides*, possessed larger AA3 and AA7 repertoires than the sampled *Bisifusarium* cheese-associated species. These families include oxidoreductases involved in carbohydrate processing, and biochemical studies have demonstrated diverse substrate preferences and interactions with polysaccharide-degrading enzymes (Haddad Momeni et al., 2021; Sützl et al., 2018). *Longinectria* also contained larger GH18 repertoires than cheese-associated *Bisifusarium*, as well as GH75 proteins absent from all sampled *Bisifusarium* genomes. These families include chitinases and chitosanases, respectively. Experimental evidence links some GH18 enzymes to antagonistic interactions (Tzelepis et al., 2015), making interactions with other cheese-associated fungi, or simply roles in the fungi’s own cell wall biology plausible. Despite these contrasts, both cheese-associated lineages lacked GH54 and GH67 present in the wild *B. dimerum* lineage and had fewer CE1 proteins. This is compatible with reduced representation of some plant polysaccharide processing functions, consistent with broader associations between CAZyme repertoires and nutritional ecology (Zhao et al., 2013). However, retention of GH74 proteins in *B. domesticum* shows that the smaller repertoire does not represent wholesale loss of plant-substrate associated functions. In cheese, complex plant polysaccharides are largely absent, and these patterns suggest selective remodelling of carbohydrate-processing repertoires.

Consistent with this functional contraction, cheese-associated *Bisifusarium* also exhibited reduced genome sizes relative to their non-cheese associated relatives. Genome reduction has been associated with ecological specialisation in fungi, where adaptation to a stable niche can relax selection on functions required in more heterogeneous environments. For example, specialised termite-associated *Podaxis* lineages have smaller genomes and reduced genome repertoires relative to free-living relatives, with losses concentrated in functions associated with their ancestral ecology (Conlon et al., 2021). Similar signatures have emerged during fungal domestication, in *Saccharomyces cerevisiae* (Legras et al., 2018) and *Geotrichum candidum* (Bennetot et al. 2023). Genome streamlining has also recently been reported during mushroom domestication, although reductions in genome size were predominantly driven by intron loss (Pérez et al., 2026). While domestication does not necessarily predict small fungal genomes, with gene or region loss without an accompanying reduction in assembly size observed in *Aspergillus oryzae* and *Penicillium* (Chang et al., 2015; Cheeseman et al., 2014; O’Donnell et al., 2025), these observations suggest that the smaller genomes of cheese-associated *Bisifusarium* may represent a broader process of genomic streamlining accompanying adaptation to a specialised anthropogenic niche.

SNP analyses revealed that, surprisingly, four out of five strains belonged to a single clonal lineage, with fewer than 400 SNPs identified among them, despite originating from different types of cheeses, produced in three different countries (Canada, France and Switzerland). Such low level of genetic variation is comparable to that observed in other domesticated cheese *Penicillium* species, such as *P. camemberti* varieties (Ropars et al., 2020), *P. biforme cheesy* (O’Donnell et al., 2026) and the non-Roquefort *P. roqueforti* cheese population (Dumas et al., 2020). These domesticated species have indeed undergone strong genetic bottlenecks due to intense selection by the cheese industry. Even more surprisingly, the known commercial culture (UBOCC-A-120004) proved to be distinct from the four other strains, constituting a distinct clonal lineage, with approximately 8,000 SNPs. This strongly suggests that at least two strains of *B. domesticum* are commercially available, and that strong genetic bottlenecks occurred as a result of intense selection. Greater diversity may exist within this species, either in cheeses or in other environments.

We also found evidence for recent horizontal gene transfer between two cheese-associated *Bisifusarium* species. A 75kb-region shared between *B. allantoides* and *B. penicillioides* exhibited >98% nucleotide identity across coding and non-coding sequence, contrasting with the 84% average nucleotide identity between the two species, strongly supporting recent horizontal gene acquisition. Several genes within this region, including ENA2, TRK, chromate transporters and phosphate transporters, correspond to the salt-tolerance gene cluster previously identified within a *Starship*-like region of cheese-associated *Penicillium* species (O’Donnell et al., 2025). Although fragmented genome assemblies prevented identification of complete *Starship* boundaries or an associated captain gene, the presence of a neighbouring SANT/Myb gene and the conservation of multiple salt-tolerance genes suggest that this region may represent a *Starship*. These findings therefore extend evidence for the role of horizontal transfer in adaptation to cheese beyond *Penicillium* fungi (Lo et al., 2023; O’Donnell et al., 2026, 2025), suggesting that exchange of stress-adaptation modules may represent a more general mechanism facilitating fungal adaptation to new environments. The fragmented short-read assemblies used in this study prevented the identification of *Starships* in cheese-associated *Nectriaceae* species, though the identification of several captains highly suggest their presence. *Starships* have been identified in other Nectriaceae species, such as the coffee pathogen *Fusarium xylarioides* (Peck et al., 2024), as well as in multiple plant pathogens of the *Fusarium* genus (Sato et al., 2025). Horizontal transfer of these *Starships* has been identified between *Fusarium* species and the distantly related pathogen *Verticillium dahliae,* which belongs to a distinct fungal family. Long-read sequencing is now needed to accurately identify *Starship* content in the cheese-associated Nectriaceae family.

An important limitation of the present study is that these fungi remain extremely poorly sampled. The isolates analysed here were recovered during a survey of only 56 artisanal cheeses, from which relatively few fusaroid strains were obtained (Savary, 2021). Beyond these isolates, very few cheese-associated *Bisifusarium* or *Longinectria* strains are publicly available. Consequently, the evolutionary patterns reported here should be regarded as hypotheses for these newly recognised cheese lineages rather than comprehensive descriptions of their diversity. Broader sampling across cheese varieties and production systems will be necessary to determine how widespread these genomic signatures are.

In conclusion, we report two remarkably different genomic trajectories for two distantly related fungal lineages in independently colonising the cheese environment. Whereas *Longinectria* primarily adapted through expansion and duplication of gene families associated with nutrient transport, regulatory and specialised metabolism, cheese-associated *Bisifusarium* followed an opposing path of genome reduction affecting many of the same functional systems. These long-term changes in genome content were accompanied by adaptive protein evolution and, at a much shorter evolutionary timescale, recent horizontal acquisition of a putative salt-tolerance cluster between *B. allantoides* and *B. penicillioides.* Interestingly, this cluster was not observed in *B. domesticum*, despite being the only known *Bisifusarium* cheese species. These results suggest that adaptation to anthropogenic environments is not constrained to a single genomic route, but can emerge through fundamentally different combinations of gene gain, gene loss and horizontal gene transfer. Yet, the ecological significance of these fungi within cheese ecosystems remains largely unknown. Several species examined here were first identified through surveys of cheese-rind diversity, and their roles in ripening, flavour development, microbial community interactions, or whether they are environmental colonists remains poorly understood. Connecting their genomic differences to substrate utilisation, salt tolerance and interactions with other cheese microorganisms will help establish their ecological roles and potential industrial importance.

## Author contributions

LP: Writing – original draft, Investigation, Formal analysis, JLJ: Writing – review and editing, Investigation, Formal analysis, MC: Writing – review and editing, OS: Investigation, Formal analysis, JPV: Formal analysis, AS: DNA extraction, TG: Funding acquisition, Writing, EC : Writing – review and editing, Supervision, Resources, Methodology, Funding acquisition, Conceptualization, JR: Writing – original draft review and editing, Supervision, Resources, Methodology, Investigation, Formal analysis, Funding acquisition, Conceptualization.

## Acknowledgements

We thank Robert Proctor (US Department of Agriculture) for sharing raw reads of *Bisifusarium domesticum* strain NRRL 29976. We thank all people who kindly sent us the various cheeses analyzed in this study (Pr. Luca Cocolin, Pr. Jérôme Mounier, Dr. Guillaume Dubrulle, Mr Sten Marc, Arnaud Lamassiaude) as well as those who have provided isolates from their research studies (Dr. Miguel Alvarez, Dr. Baltasar Mayo and Franck Deniel). We thank the UBO Culture collection (https://www.univbrest.fr/ubocc/fr) for strains preservation and distribution.

