## Supplementary Figures for "Cheese adaptation beyond *Penicillium*: contrasting genomic trajectories in the distantly related *Nectriaceae* molds"

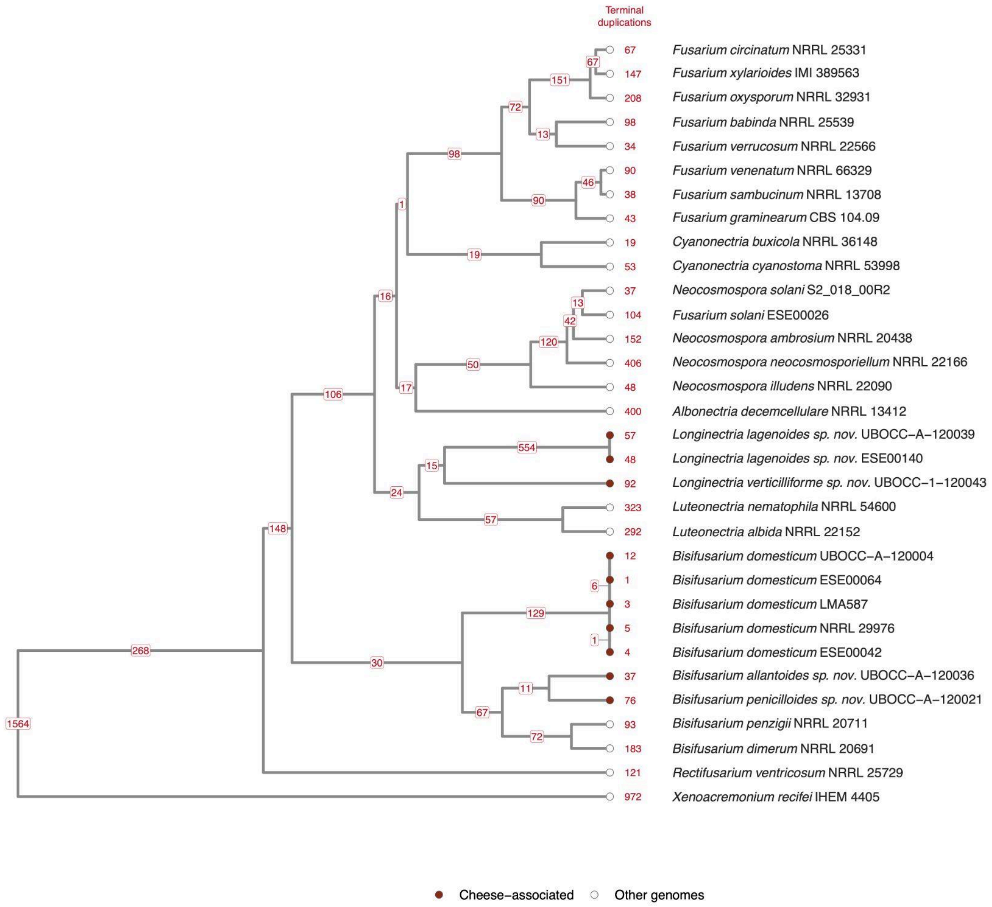


Figure S1. *Nectriaceae* species tree with OrthoFinder gene duplication events mapped to branches. Red labels on internal branches give retained duplication counts and the column gives counts on terminal branches. Both copies must occur in at least 50% of descendant genomes, and in both genomes for branches subtending two genomes. Terminal events carrying assembly/ annotation artefact flags are excluded. Branch lengths follow the input ultrametric tree.

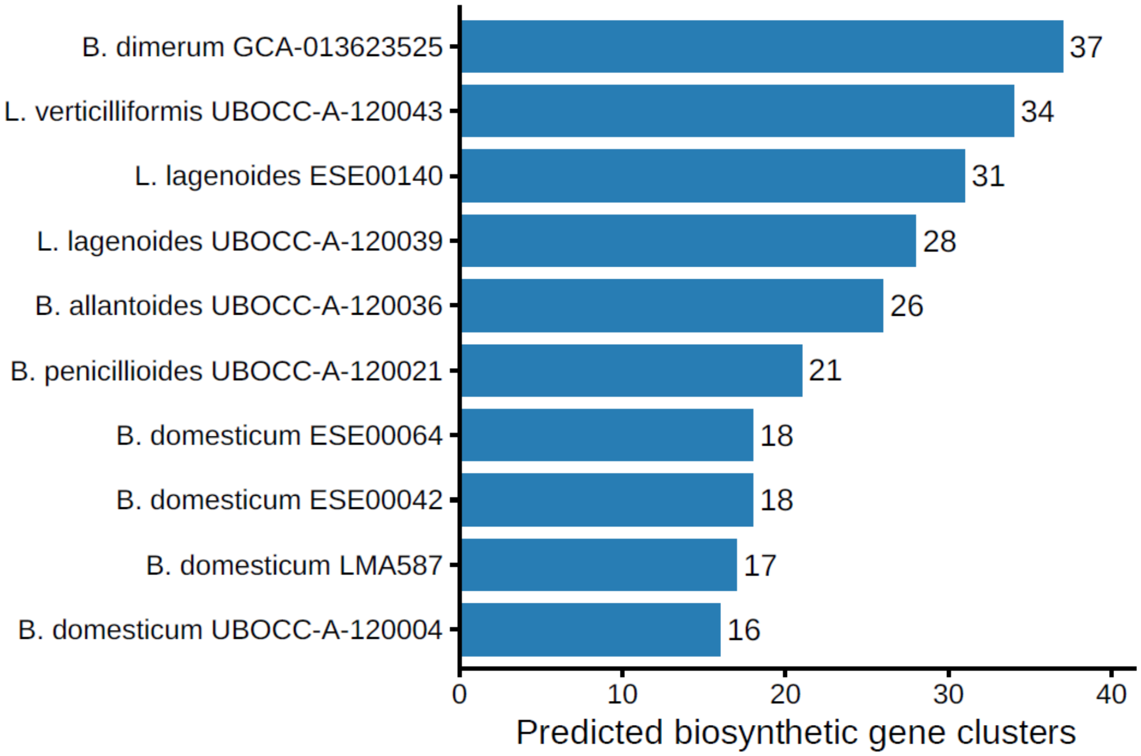


Figure S2. Number of predicted biosynthetic gene clusters identified in each genome. Bars indicate the total number of biosynthetic gene clusters predicted by antiSMASH v8.0 for each annotated genome. Genome fragmentation may result in partial or split BGC predictions.

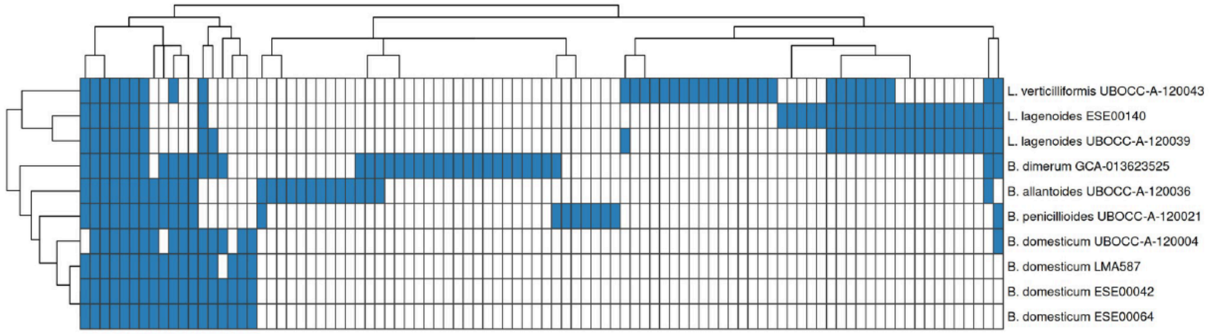


Figure S3. Presence/absence of Gene Cluster Families across genomes. Columns represent BiG-SCAPE Gene Cluster Families, and rows represent genomes. Blue cells indicate the presence of a biosynthetic gene cluster belonging to a given gene cluster family, whereas white cells indicate its absence. Hierarchical clustering was performed on both genomes and gene cluster families using Euclidean distances calculated from the binary presence/absence matrix and complete linkage, with the resulting dendrograms shown on the left and above the matrix, respectively. Because some genomes are fragmented, the absence of a gene cluster family should be interpreted with caution, as incomplete genome assemblies may lead to missing or fragmented biosynthetic gene cluster predictions.

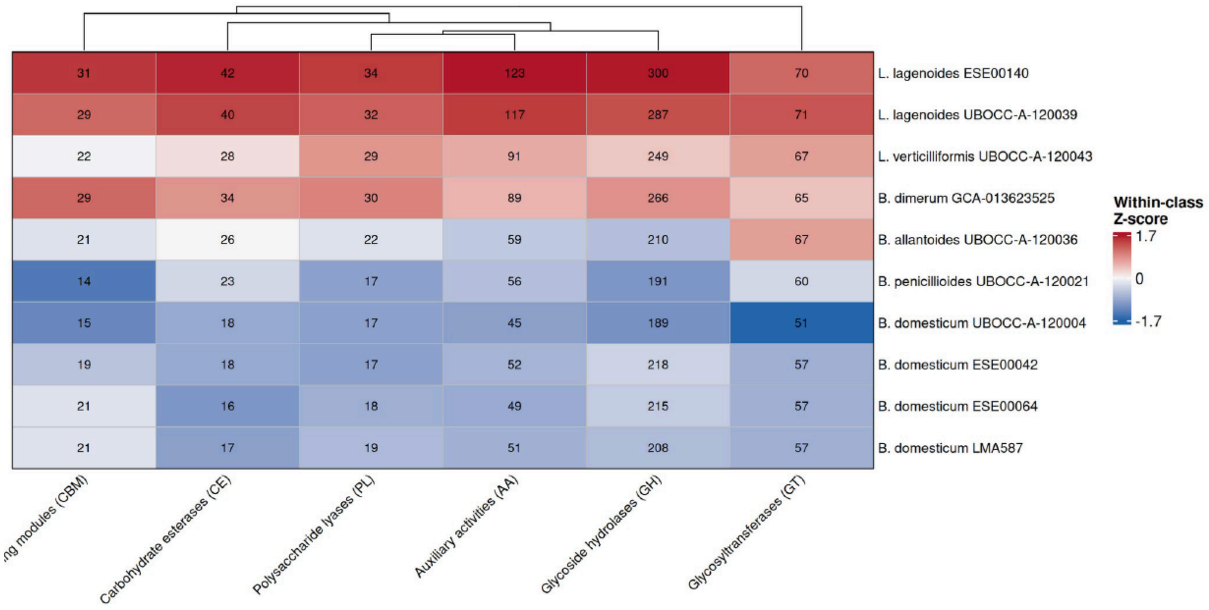


Figure S4. Distribution of CAZy classes across genomes. Numbers indicate the raw counts of CAZy annotations assigned to each functional class. Cell colors represent Z-scores calculated independently for each CAZy class across genomes, with red and blue indicating values above and below the class mean, respectively. Columns were hierarchically clustered based on their standardized abundance profiles.

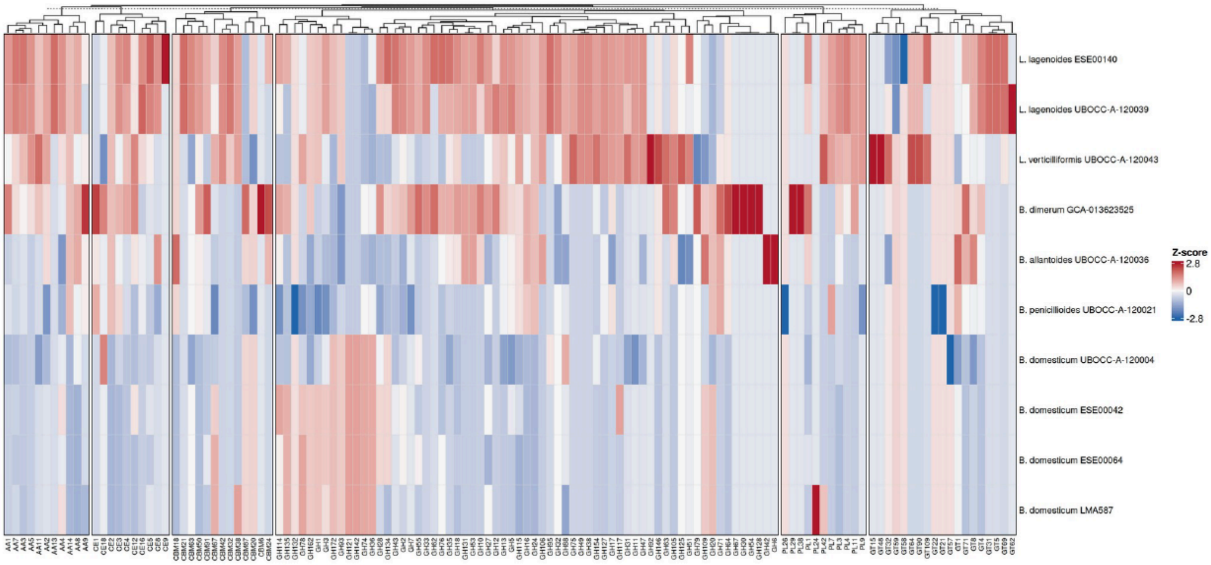


Figure S5. Comparative distribution of CAZy families across genomes. Columns represent individual CAZy families grouped according to their major functional class, and rows represent genomes. For each family, annotation counts were standardized across genomes as Z-scores; red and blue therefore indicate relative enrichment and depletion, respectively, compared with the mean abundance of that family. CAZy families were hierarchically clustered within each functional class.

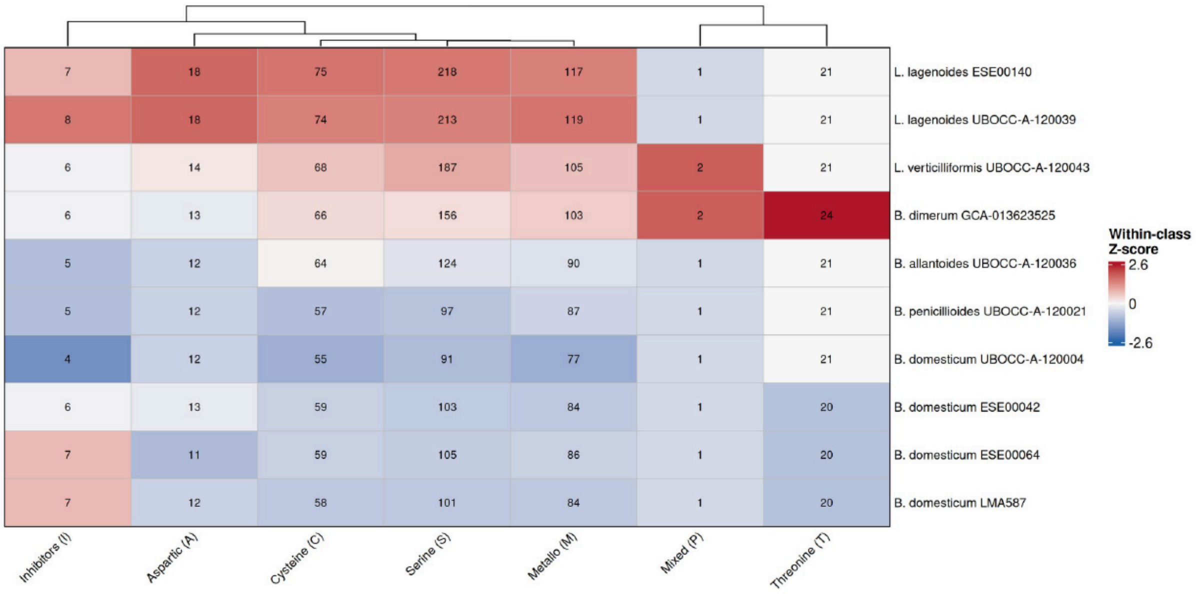


Figure S6. Distribution of microbial eukaryote reference oriented peptidase (MEROPs) peptidase classes across genomes. Numbers indicate the raw counts of peptidase annotations assigned to each MEROPS catalytic class. Cell colors represent Z-scores calculated independently for each class across genomes, with red and blue indicating values above and below the class mean, respectively. Columns were hierarchically clustered based on their standardized abundance profiles..

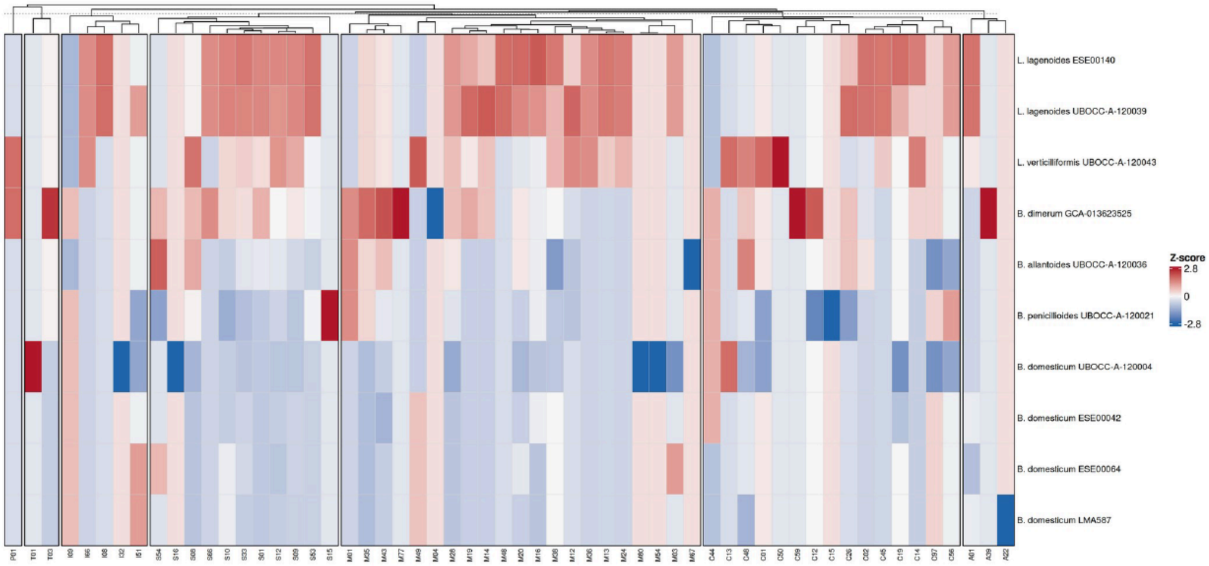


Figure S7. Comparative distribution of microbial eukaryote reference oriented peptidase (MEROPs) peptidase families across genomes. Columns represent individual MEROPS families grouped according to their catalytic class, and rows represent genomes. For each family, annotation counts were standardized across genomes as Z-scores; red and blue indicate relative enrichment and depletion, respectively, compared with the mean abundance of that family. Peptidase families were hierarchically clustered within each catalytic class.

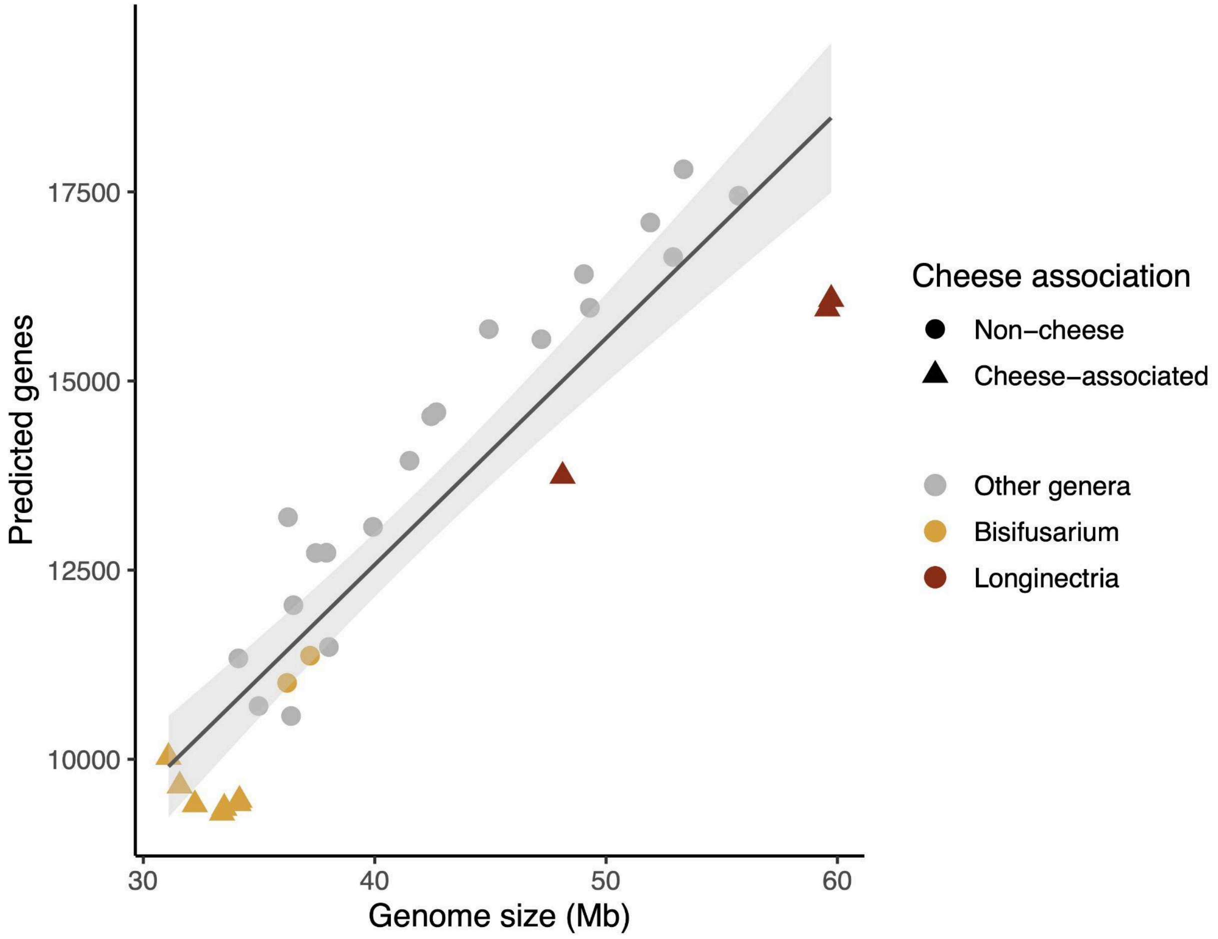


Figure S8. Relationship between genome size and predicted gene number. Each point represents a genome, coloured by genus: *Bisifusarium* (yellow); *Longinectria* (red); all other genera (grey). Triangles indicate cheese-associated genomes, and circles indicate non-cheese genomes. Solid line shows the linear regression fitted across all genomes, with the shaded band indicating 95% confidence interval.

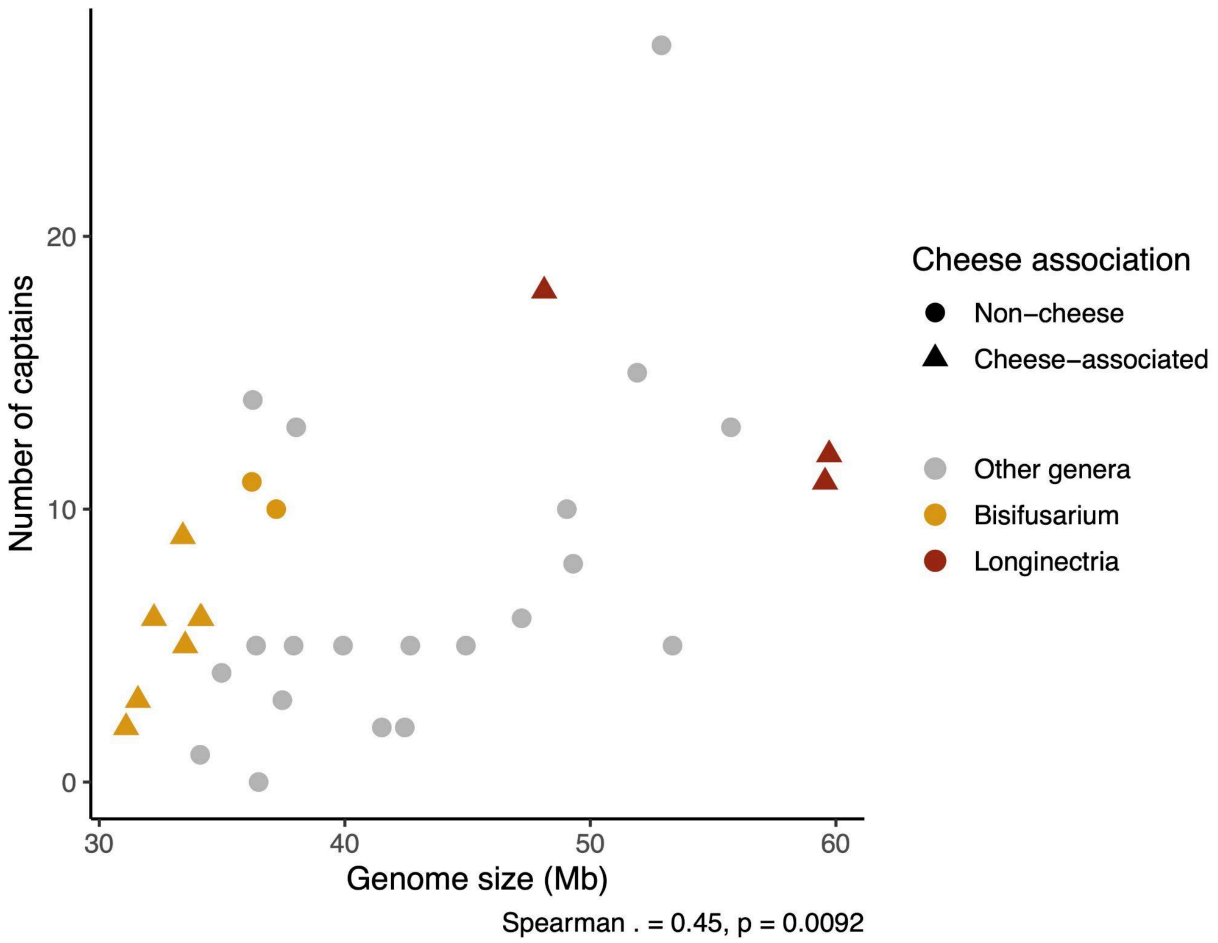


Figure S9. Relationship between genome size and predicted number of *Starship* captains. Each point represents a genome, coloured by genus: *Bisifusarium* (yellow); *Longinectria* (red); all other genera (grey). Triangles indicate cheese-associated genomes, and circles indicate non-cheese genomes. Genome size was positively associated with captain number (Spearman's rho = 0.45; *P* = 0.0092).
